# Nsp1 of SARS-CoV-2 Stimulates Host Translation Termination

**DOI:** 10.1101/2020.11.11.377739

**Authors:** Alexey Shuvalov, Ekaterina Shuvalova, Nikita Biziaev, Elizaveta Sokolova, Konstantin Evmenov, Tatiana Egorova, Elena Alkalaeva

## Abstract

The Nsp1 protein of SARS-CoV-2 regulates the translation of host and viral mRNAs in cells. Nsp1 inhibits host translation initiation by occluding the entry channel of the 40S ribosome subunit. The structural study of SARS-CoV-2 Nsp1-ribosomal complexes reported post-termination 80S complex containing Nsp1 and the eRF1 and ABCE1 proteins. Considering the presence of Nsp1 in the post-termination 80S ribosomal complex simultaneously with eRF1, we hypothesized that Nsp1 may be involved in translation termination. Using a cell-free translation system and reconstituted *in vitro* translation system, we show that Nsp1 stimulates translation termination in the stop codon recognition stage at all three stop codons. This stimulation targets the release factor 1 (eRF1) and does not affect the release factor 3 (eRF3). The activity of Nsp1 in translation termination is provided by its N-terminal domain and the minimal required part of eRF1 is NM domain. We assume that biological meaning of Nsp1 activity in translation termination is binding with the 80S ribosomes translating host mRNAs and removal them from the pool of the active ribosomes.

## INTRODUCTION

Currently, the new coronavirus infection COVID-19 pandemic continues around the world, having already killed more than 1 million people. The causative agent of COVID-19 is Severe Acute Respiratory Syndrome Coronavirus 2 (SARS-CoV-2), which belongs to the beta-coronavirus group (Zhu et al., 2020). Its genome consists of a single-stranded (+) RNA, approximately 30,000 nt long, containing 14 open reading frames (ORFs) encoding 27 proteins. The two overlapping 5’-end frames ORF1a and ORF1b encode large polypeptides (490 and 794 kDa, respectively), which are subsequently proteolytically cleaved by viral proteases (Nsp3 and Nsp5), generating the unstructured proteins Nsp1-Nsp16. The ORF1b polypeptide synthesis occurs via a −1 frameshift just before the ORF1a stop codon. The RNA-dependent RNA polymerase NSP12 encoded within the ORF1b ensures the replication of the viral genome, also producing additional truncated RNA products with open reading frames for the synthesis of the remaining SARS-CoV-2 proteins (Kim et al., 2020b; Wu et al., 2020; Zhou et al., 2020). It has been shown that among all SARS-CoV-2 proteins, Nsp1 is a major viral factor that affects cellular viability (Yuan et al., 2020).

Nsp1 is the first protein produced by a virus. It affects the vital activity of the infected cell and is necessary for the development of viral particles. Nsp1 induces apoptosis, and alters the transcriptional profile of cells by blocking the transcription of genes responsible for cell metabolism, regulation of the cell cycle, mitochondrial function, antigen presentation, ubiquitin/proteasome pathways, and protein synthesis (Yuan et al., 2020). Moreover, Nsp1 directly inhibits translation both *in vitro* in the cell-free translation systems and *in vivo* in the cells (Banerjee et al., 2020; Schubert et al., 2020; Shi et al., 2020; Tidu et al., 2020; Yuan et al., 2020). Therewith, viral mRNAs are less susceptible to translation inhibition due to the presence of an SL1 hairpin in their 5’UTR (Banerjee et al., 2020; Shi et al., 2020; Tidu et al., 2020). Nsp1-mediated inhibition of translation leads to suppression of the host interferon response, which is one of the main mechanisms of cellular defense against viral replication (Banerjee et al., 2020; Thoms et al., 2020; Xia et al., 2020).

Nsp1 of the SARS-CoV-2 virus is a small protein of 180 amino acid residues, consisting of N- and C-terminal domains connected by an unstructured linker (Shi et al., 2020; Wu et al., 2020). The N-domain of this protein interacts with the 5’UTR of SARS-CoV-2 mRNA, which is necessary for its efficient translation (Shi et al., 2020). Mutations in the N-domain also lead to a decrease in the level of apoptosis of infected cells (Yuan et al., 2020). The C-domain of Nsp1 binds to the entry channel of the 40S ribosome subunit. In this case, the 40S subunit associated with Nsp1 binds to mRNA much worse and is unable to form a native translation initiation complex. Thus, Nsp1 inhibits the initiation of host mRNA translation (Banerjee et al., 2020; Lapointe et al., 2020; Schubert et al., 2020; Shi et al., 2020; Thoms et al., 2020; Tidu et al., 2020; Yuan et al., 2020). In addition to free 40S subunits, Nsp1 SARS-CoV-2 is also found in aberrant 48S initiator complexes and 80S ribosomes (Lapointe et al., 2020; Schubert et al., 2020; Thoms et al., 2020; Yuan et al., 2020). It should be noted that binding to the 40S subunit of homologous Nsp1 in closely related SARS-CoV does not block the formation of the 48S initiator complex but inhibits the attachment of the 60S subunit to it and causes inhibition of the translation initiation (Kamitani et al., 2009). Some of the 80S ribosomes with Nsp1 SARS-CoV-2 contain additional proteins involved in translation termination and ribosome recycling, i.e., the eukaryotic release factor eRF1 and ABCE1 (Thoms et al., 2020). This prompted us to address a possible role of Nsp1 in translation termination.

Translation termination is the final stage of protein biosynthesis, in which the nascent polypeptide releases from the ribosome. Translation termination begins when one of the three stop codons appears in the A site of the ribosome (Jackson et al., 2012). The ribosome then binds eRF1 that recognizes all three stop codons (Brown et al., 2015; Frolova et al., 1994; Kryuchkova et al., 2013; Matheisl et al., 2015; Song et al., 2000). A partner protein of eRF1 is eRF3, the GTPase that activates eRF1 (Alkalaeva et al., 2006; Cheng et al., 2009; Frolova et al., 1996; Shao et al., 2016; Taylor et al., 2012). Concerted binding of the eRF1-eRF3-GTP complex with the ribosome stimulates GTPase activity of eRF3, which, in turn, causes a conformational change of eRF1 (Cheng et al., 2009). As a result, the catalytic GGQ motif of eRF1 is juxtaposed to the peptidyl transferase center of the ribosome. This induces the hydrolysis of peptidyl-tRNA and the release of the synthesized protein (Alkalaeva et al., 2006; Brown et al., 2015; Frolova et al., 1999; Jackson et al., 2012; Matheisl et al., 2015; Song et al., 2000). Termination of translation is one of the critical stages of protein biosynthesis because its suppression allows for inhibiting both the release of peptides from the ribosomes and the transition of the ribosomes to recycling, consequently, to prevent new rounds of translation.

The simultaneous binding of the eRF1 and Nsp1 to the 80S ribosomes (Thoms et al., 2020), implies a possible role of Nsp1 in translation termination, which has not been elucidated yet. To study an effect of Nsp1 on the termination of host translation, we used the reconstituted mammalian translation system (Alkalaeva et al., 2006) as well as pre-termination complexes (preTCs) purified from rabbit reticulocyte lysate (RRL) (Susorov et al., 2020). We demonstrate that Nsp1 of SARS-CoV-2 stimulates translation termination and determines the stage of termination at which it works. Direct involvement in translation termination of Nsp1 was confirmed by studying activities of the mutants of Nsp1. The data obtained allow us to propose a possible mechanism of control of translation on the stage of termination by SARS-CoV-2 Nsp1.

## RESULTS

### Nsp1 stimulates peptide release

To study the activity of Nsp1 in translation termination, the codon-optimized sequence of the SARS-CoV-2 Nsp1 was cloned into the petSUMO vector and expressed in *E. coli* BL21(DE3) cells. Noteworthy, the resulting Nsp1 lacked any tag. We first verified the ability of the recombinant Nsp1 to inhibit translation. For this purpose, translation of Nluc mRNA in RRL was assayed in the presence of Nsp1 (Fig. 1A). We confirmed that our preparation of the recombinant tag-free Nsp1 significantly inhibited the translation of Nluc mRNA in the cell-free system.

**Figure 1.**
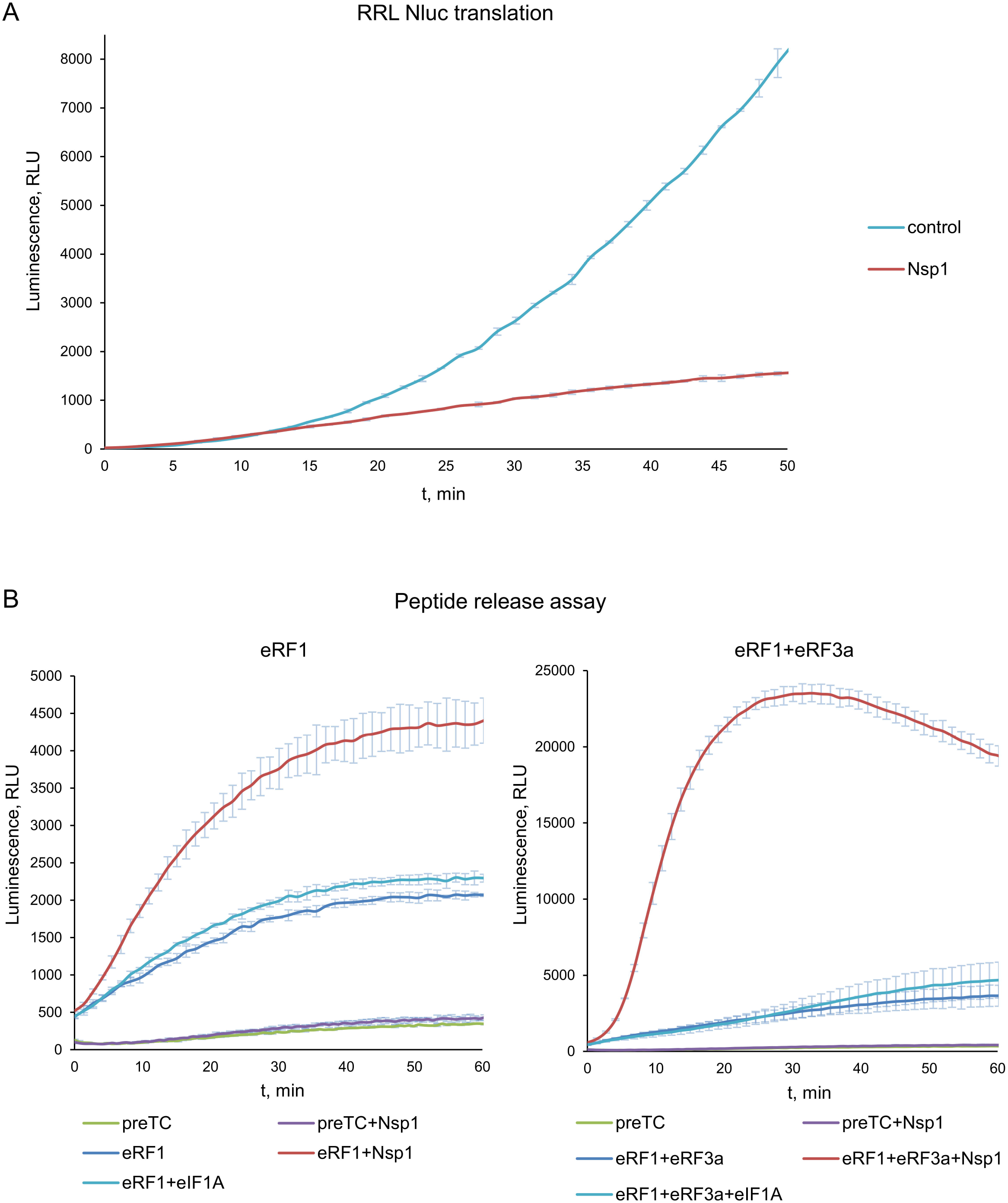
Nsp1 activates peptidyl-tRNA hydrolysis with the release factors. (A) *In vitro* Nluc mRNA translation in RRL in presence/absence of Nsp1. Time progress curves showing luminescence (in relative luminescence units, rlu), n=3, mean±SD (B) Termi-Luc peptide release assay in presence/absence of Nsp1 and the release factors. Time progress curves showing luminescence (in relative luminescence units, rlu) with NLuc released from ribosome complex upon treatment with the proteins of interest. eIF1A was added to the reaction as a negative control. n=3, mean±SD.

Next, we evaluated a possible effect of SARS-CoV-2 Nsp1 on translation termination. The result of translation termination is the hydrolysis of peptidyl-tRNA and the release of the nascent peptide from the ribosome. We performed an analysis of a peptide release efficiency using the Termi-Luc approach developed recently by Susorov et al. (2020). Pretermination complexes PreTCs, containing NanoLuc luciferase (Nluc) located at the P site of the ribosome in the peptidyl-tRNA form and UAA stop codon at the A site, were purified from the RRL. The addition of the release factors to the purified preTC triggers the peptidyl-tRNA hydrolysis and induces the release of Nluc from the ribosome. Human release factors eRF1 and eRF3a, together with Nsp1, were added to the preTC, and the appearance of luminescence was measured over time. Nsp1 is unable to perform peptide release by itself, however it significantly stimulates translation termination induced by eRF1 alone and by the complex of eRF1-eRF3a (Fig. 1B). As a negative control we used eukaryotic initiation translation factor eIF1A, which is of a similar size and binds close to the Nsp1 binding site on the ribosome. Interestingly, eIF1A was shown to be involved into ribosomal recycling after translation termination (Pisarev et al., 2007), however here we did not detect its influence on peptide release. Probably it was due to the absence in the mixture of the other eukaryotic initiation factors and the recycling protein ABCE1.

Considering data on Nsp1 activity in peptide release, we concluded that the presence of Nsp1 in the 80S ribosomal complexes (Thoms et al., 2020) reflects its functional significance in translation termination.

### Nsp1 stimulates the termination complexes formation

To study a molecular mechanism of the stimulation of translation termination by Nsp1, we tested its activity on the purified preTCs obtained in the reconstituted translation system (Alkalaeva et al., 2006). We assembled preTCs from individual components on the model MVHL mRNAs with the UAA stop codon and purified them by centrifugation in a sucrose density gradient. These complexes were then used to test the effect of Nsp1 on the formation of termination complexes (TCs) by fluorescent toe-print analysis, which shows the position of ribosomal complexes on mRNA. When stop codon is recognized by eRF1, the ribosome changes conformation of the +4 nucleotide following the stop codon, which is then detected as a nucleotide shift of the toe-print signal (Alkalaeva et al., 2006; Egorova et al., 2019). It should be noted that in each experiment we compared the relative amounts of cDNAs as areas of the preTC and the TC peaks according to the following formula TC formation efficiency = TC / (TC + preTC), which is presented on the histograms (Fig. 2).

**Figure 2.**
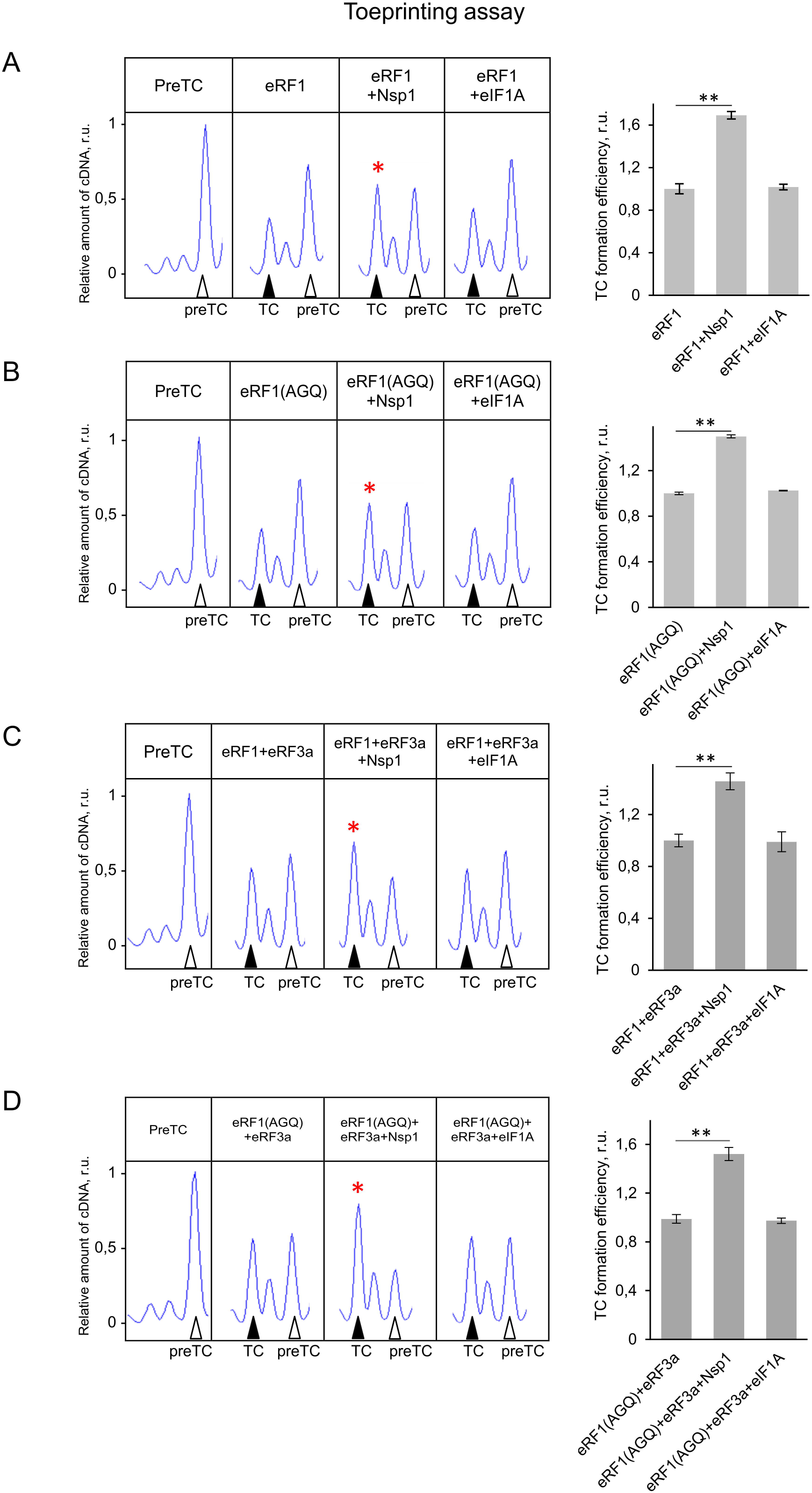
Nsp1 increases TC formation. Examples of raw toe-printing data and relative quantitative analysis of the TC formation efficiency induced by the release factors in the presence of Nsp1 or eIF1A as a negative control. TC formation efficiency induced by release factors was taken as 1. (A) Toe-print analysis of termination complexes formed by addition to the preTCs of eRF1 and Nsp1 or eIF1A as a negative control. (B) Toe-print analysis of termination complexes formed by addition to the preTCs of eRF1(AGQ) and Nsp1 or eIF1A as a negative control. (C) Toe-print analysis of termination complexes formed by addition to the preTCs of eRF1+eRF3a+GTP and Nsp1 or eIF1A as a negative control. (D) Toe-print analysis of termination complexes formed by addition to the preTCs of eRF1(AGQ)+eRF3a+GTP and Nsp1 or eIF1A as a negative control. r.u. – relative units. Positions of preTCs and TCs are labeled by white and black triangles, respectively. Red stars indicate the increased quantity of ribosomal complexes, shifted from the preTC to the TC state. The error bars represent the standard error of mean, stars (**) mark a significant difference from the respective control *P* < 0.01 (n=3).

Toe-print analysis showed that the addition of Nsp1 to the preTC does not induce any shift of the preTC peak both in 0,5 and 2,5 μM concentration (Fig. S1A). However, addition of Nsp1 to the preTC with the release factor eRF1 increases the amount of the formed TC by 1,6 times (Fig. 2A). As a negative control, we also used eIF1A. To determine the stage of termination during which Nsp1 activates TC formation, we tested the activity of the eRF1(AGQ) mutant, which is unable to induce hydrolysis of peptidyl-tRNA. Efficiency of the TC formation in the presence of AGQ mutant reflects the efficiency of eRF1 binding with the stop codon irrespective of peptidyl-tRNA hydrolysis. We observed that TC formation performed by eRF1(AGQ) was similarly stimulated by Nsp1 (Fig. 2B).

Additionally, Nsp1 stimulates the formation of the TC in the presence of both release factors (eRF1 and eRF3a) and GTP (Fig. 2C). Likewise, in the presence of the eRF1(AGQ)-eRF3a complex, we observed significant stimulation of TC formation by Nsp1 (Fig. 2D). Thus, we concluded that Nsp1 stimulates translation termination before peptide release either during the binding of eRF1 with the stop codon or during GTP hydrolysis catalyzed by eRF3a.

### Nsp1 does not influence the GTPase activity of eRF3a

One of the stages of translation termination is the GTP hydrolysis performed by eRF3, which occurs after the binding of eRF1-eRF3 complex to the stop codon inside the ribosome. This is followed by a conformational change of the TC and positioning of eRF1 in the peptidyl-transferase center of the ribosome (Cheng et al., 2009). By analyzing the GTPase activity of eRF3a in the presence of eRF1, 80S ribosome, and Nsp1, we demonstrated no effect of Nsp1 on the hydrolysis of GTP (Fig. 3A). The GTPase activity of eRF3a was determined at the concentrations of the components corresponding to the linear part of the saturation curve (Fig. S1B). We confirmed this finding by analyzing the efficiency of TC formation in the presence of a non-hydrolysable GTP analogue, GDPCP, using toe-printing assay. The addition of the GDPCP to eRF1 and eRF3a did not prevent stimulation of TC formation by Nsp1 similar to that observed in the presence of GTP (Fig. 3B). Therefore, Nsp1 stimulation activity in translation termination appears before and independently of GTP hydrolysis catalyzed by eRF3.

**Figure 3.**
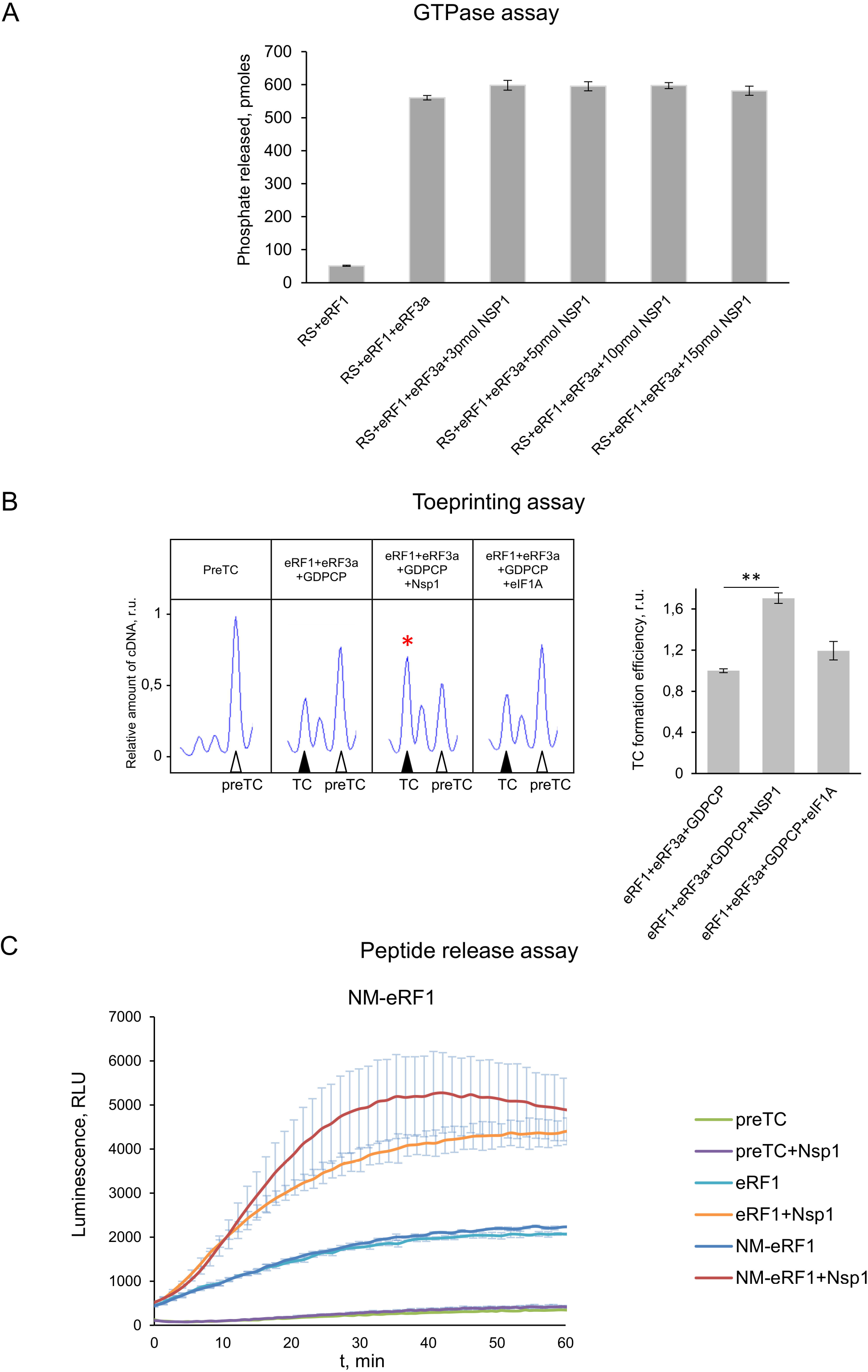
Nsp1 stimulates translation termination independently of GTP hydrolysis. (A) GTPase activity of the eRF3a in the presence of the ribosome subunits, eRF1 and Nsp1, mean±SE, (n=4). (B) Nsp1 stimulation of TC formation does not depend on GTP hydrolysis. TC formation was induced by the addition of eRF1+eRF3a+GDPCP and Nsp1 or eIF1A as a negative control. An example of raw toe-printing data and relative quantitative analysis of the TC formation efficiency by eRF1+eRF3a+GDPCP in the presence of Nsp1 or eIF1A as a negative control. TC formation efficiency induced by eRF1+eRF3a+GDPCP was taken as 1. The TC corresponds to the black triangle, and the preTC corresponds to the white triangle, r.u. – relative units. Red star indicates the increased quantity of ribosomal complexes, shifted from the preTC to the TC state. The error bars represent the standard error of mean, stars (**) mark a significant difference from the respective control *P* < 0.01 (n=3). (C) Termi-Luc peptide release assay in the presence/absence of Nsp1 and NM-eRF1 or eRF1. Time progress curves showing luminescence (in relative luminescence units, rlu) with NLuc released from the ribosome complex upon treatment with the proteins of interest, n=3, mean±SD.

Additionally, we tested the ability of Nsp1 to stimulate peptide release induced by truncated from the C-end form of eRF1, NM-eRF1, which is unable to interact with eRF3a. We observed even the higher level of peptide release stimulation of NM-eRF1 by Nsp1 (Fig. 3C). This once again confirms our conclusion that Nsp1 does not affect the GTPase activity of eRF3, and the stimulation of translation termination by Nsp1 occurs on the stage of stop codon recognition.

### Effects of mutations of Nsp1 on its activity in translation termination

To find critical regions of Nsp1 to perform its function during termination, we obtained five mutant forms of Nsp1: K164A/H165A (KH/AA), Y154A/F157A (YF/AA), R171E/R175E (RR/EE), R124S/K125E (RK/SE), and N128S/K129E (NK/SE), described previously as proteins that lack an activity in translation or mutations that disrupt cell apoptosis (Fig. 4A) (Schubert et al., 2020; Thoms et al., 2020; Yuan et al., 2020). The mutant proteins KH/AA, YF/AA, and RR/EE were unable to suppress translation in the cell-free system (Fig. 4B), in agreement to the previous studies (Schubert et al., 2020; Thoms et al., 2020). Moreover, the mutants KH/AA and YF/AA slightly activated cell-free translation. However, the mutants RK/SE, NK/SE decreased the translation of the Nluc in cell-free system similarly to the wild-type Nsp1 (Fig. 4B). On the other hand, the mutants KH/AA and YF/AA stimulated peptide release in the presence of eRF1 alone and the eRF1-eRF3a complex similarly to Nsp1 (Fig. 4C). However, the mutants RR/EE, RK/SE and NK/SE lost the ability to stimulate peptide release (Fig. 4C).

**Figure 4.**
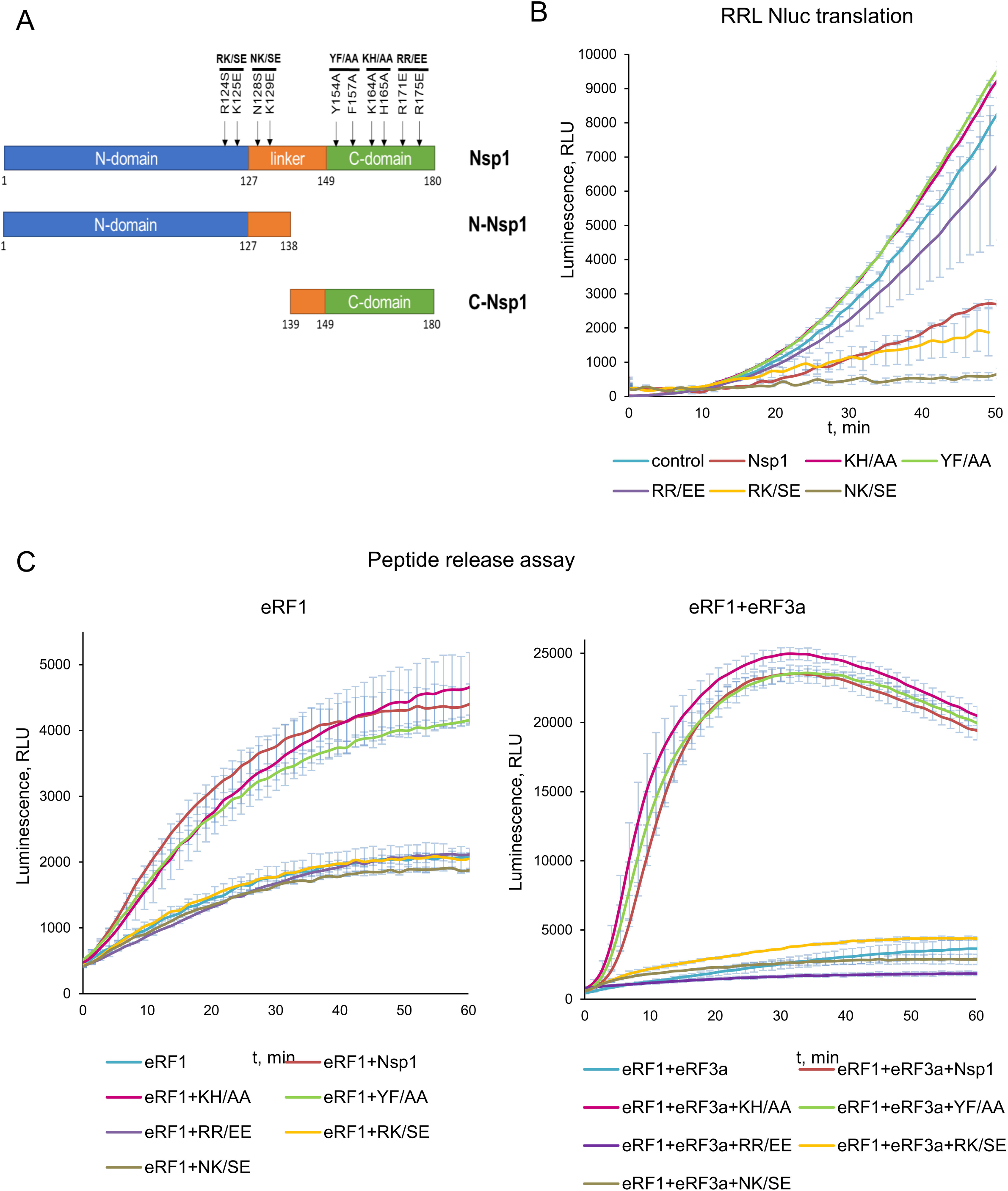
Activity of the Nsp1 mutants in translation termination. (A) Scheme of the Nsp1 constructions used in the experiments: Nsp1 N/C domains and Nsp1 mutants. (B) *In vitro* Nluc mRNA translation in RRL in the presence/absence of Nsp1 mutants. Time progress curves showing luminescence (in relative luminescence units, rlu). (C) Termi-Luc peptide release assay in the presence/absence of Nsp1 mutants and the release factors. Time progress curves showing luminescence (in relative luminescence units, rlu) with NLuc released from the ribosome complex upon treatment with the proteins of interest, n=3, mean±SD.

TC formation analysis of the same mutants showed that the proteins KH/AA and YF/AA demonstrated the same activity as Nsp1 wt (Fig. S2). On the contrary, the mutants RR/EE, RK/SE and NK/SE did not stimulate the formation of TC (Fig. S2). Therefore, TC formation analysis confirmed the results obtained by the peptide release assay.

We believe that activation of the cell-free translation (Fig. 4B) and stimulation of peptide release (Fig. 4C) by the two mutant proteins KH/AA and YF/AA reflect their positive effect on translation termination in combination with the loss of the ability to inhibit initiation. On the other hand, the mutant RR/EE was inactive in all performed tests, so we concluded that this mutation reduces the whole activity of the protein, probably disrupting its structure. Indeed, the most part of expressed RR/EE precipitated during isolation from the lysate and further purifications, which indicates a violation of its structure.

Summarizing the obtained results, we can conclude that different parts of Nsp1 are responsible for different activities of this protein in translation. The analysis of mutant activities indicated that the C domain of Nsp1 is involved in translation repression via binding with the 40S subunit, which was shown by structural analysis (Schubert et al., 2020; Thoms et al., 2020; Yuan et al., 2020), but the N domain of Nsp1 is responsible for the activation of translation termination.

We also tested 3xFLAG-tagged Nsp1 (3xFLAG-Nsp1) in peptide release and TC formation analysis, as this form of Nsp1 has been used in different structural and biochemical studies (Schubert et al., 2020; Thoms et al., 2020; Yuan et al., 2020). Interestingly, similar to untagged Nsp1 amount of 3xFLAG-Nsp1 did not suppress translation of Nluc mRNA in the cell-free system (Fig. S3A). However, in the presence of either eRF1 alone, or the eRF1-eRF3a-GTP complex, 3xFLAG-Nsp1 stimulated TC formation and peptidyl-tRNA hydrolysis, like the wild-type Nsp1 (Fig. S3B,C). Thus, the 3xFLAG does not interfere with the functioning of Nsp1 in translation termination but prevents suppression of protein translation in the cell-free system. We did not find any information about influence of 3xFLAG-Nsp1 on translation in cell-free systems. All published data describe translation activity of 3xFLAG-Nsp1 or its mutants during overexpression in cell lines. Although 3xFLAG-Nsp1 partially suppress translation in cell lines, overexpression of this protein in the cells could give inaccurate results due to the high amount of low activity protein. As far as we have found nobody compared activities in translation of untagged Nsp1 and 3xFLAG-Nsp1. Probably an extremely high negative charge from 3xFLAG-tag (at pH 7,5 charge is −7.17) interferes with the interaction of this protein with the ribosomal RNA. It in turn can cause an ineffective translational suppression and low amount of purified ribosomal complexes from cell lysates via FLAG antibodies.

### Individual N and C domains of Nsp1 are inactive in the initiation and termination of translation

It would be interesting for further research to determine whether it is possible to work with individual Nsp1 domains. Therefore, we decided to see if we can localize the activities in initiation and termination in separate domains of Nsp1. We obtained the N domain that contained 1-138 amino acid residues, and the C domain that contained 139-180 amino acid residues (Fig. 4A). Then we tested the effect of the individual Nsp1 domains on the cell-free translation and peptide release. Unfortunately, both domains lost their function in suppressing NLuc cell-free translation (Fig. 5A) and did not stimulate hydrolysis of peptidyl-tRNA in the presence of eRF1 or the eRF1-eRF3a complex (Fig. 5B). Only the N domain of Nsp1 slightly improved the peptide release in the presence of eRF1 with eRF3a (Fig. 5B).

**Figure 5.**
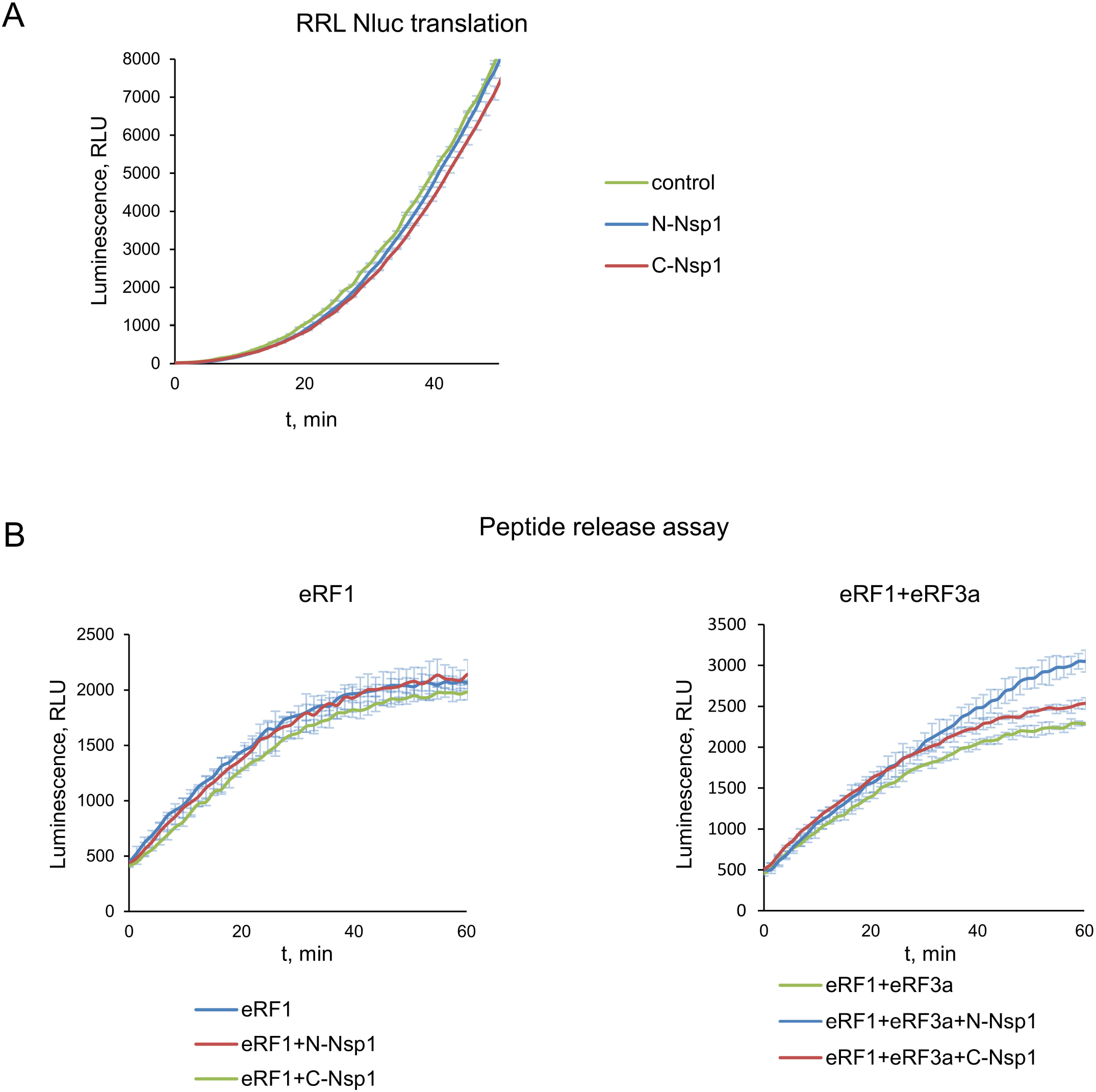
Activity of the Nsp1 domains in translation termination. (A) *In vitro* Nluc mRNA translation in RRL in the presence/absence N/C domains of Nsp1, n=3, mean±SD. (B) Termi-Luc peptide release assay in the presence/absence of N/C domains of Nsp1 and the release factors. Time progress curves showing luminescence (in relative luminescence units, rlu) with NLuc released from the ribosome complex upon treatment with the proteins of interest, n=3, mean±SD.

Activities of the N and the C domains of Nsp1 were also checked by TC formation analysis. The activities of the domains clearly follow the pattern of their activities in the peptide release. The C domain of Nsp1 was unable to stimulate TC formation, and the N domain slightly stimulated TC formation in the presence of eRF1 (Fig. S4). Thus, we have shown that the functioning of Nsp1, both in the translation termination and cell-free translation, requires the presence of the full-length protein.

### Nsp1 stimulates translation termination at all three stop codons

As we obtained results of stimulation of peptide release and TC formation by Nsp1 only at UAA stop codon, we studied its activity at other codons. First, we determined the efficiency of TC formation on the model mRNAs with UAA, UAG and UGA stop codons (Fig. 6). We revealed that in all three stop codons, Nsp1 equally stimulated TC formation in the presence of eRF1(AGQ) (Fig. 6A) and the eRF1(AGQ)-eRF3-GTP complex (Fig. 6B). Second, we tested whether in the presence of Nsp1 appears a capability of eRF1 to recognize sense codons. We determined the efficiency of TC formation on the model mRNAs with UGG and GCU codons (Fig. S5). We revealed that at the sense codons, Nsp1 was unable to induce TC formation in the presence of the eRF1(AGQ)-eRF3-GTP complex. Therefore, Nsp1 is equally effective in stimulating of translation termination on all three stop codons and does not work on the sense codons. This confirms the specific effect of Nsp1 on translation termination.

**Figure 6.**
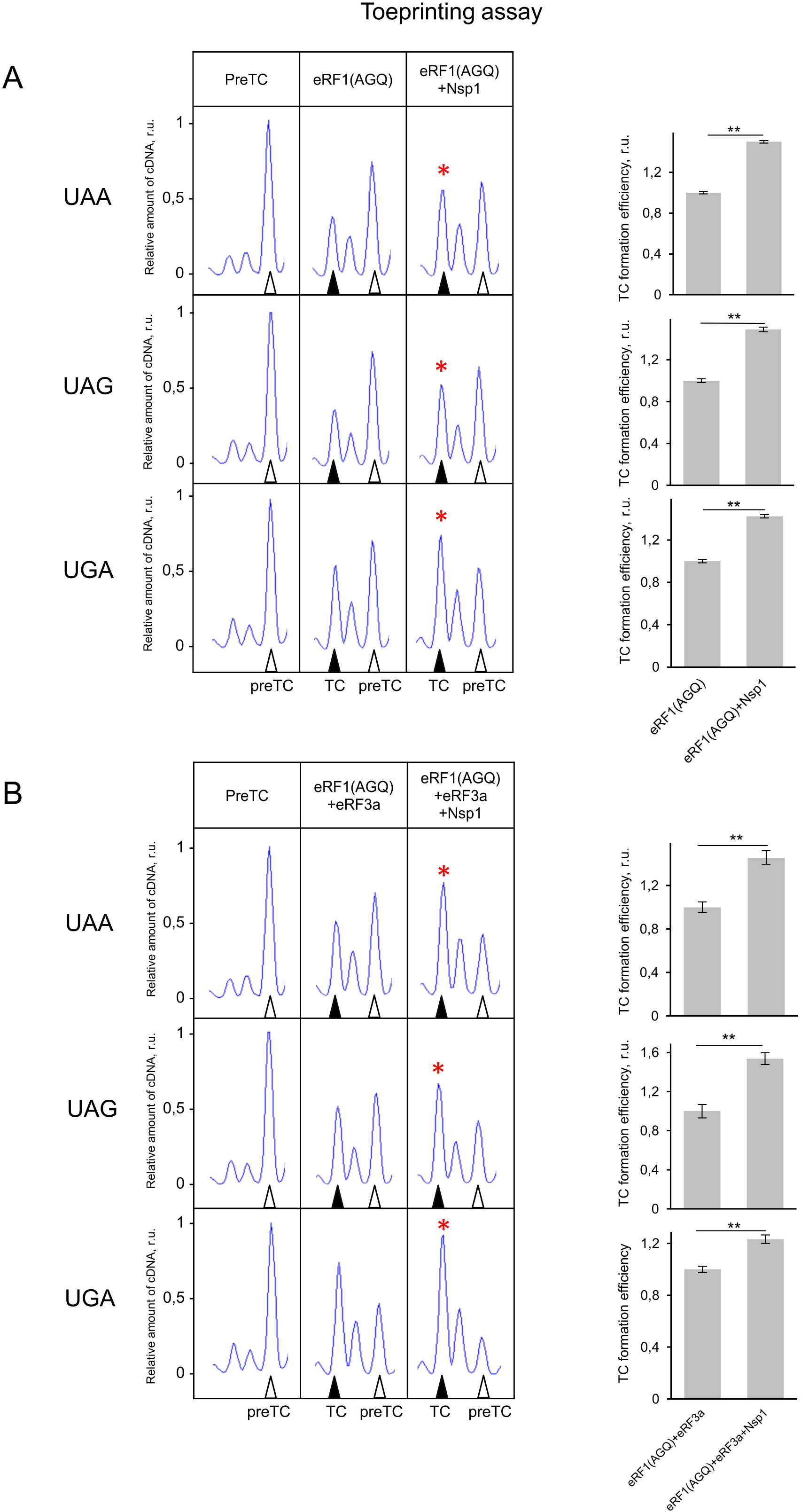
NSP1 increases TC formation on all three stop codons. (A) Toe-print analysis of termination complexes formed by addition to the preTCs of eRF1 and Nsp1 and relative quantitative analysis of the TC formation efficiency on different stop codons (UAA, UAG, UGA) in the presence of eRF1 and Nsp1. TC formation efficiency induced by eRF1 alone was taken as 1. (B) Toe-print analysis of termination complexes formed by addition to the preTCs on different stop codons (UAA, UAG, UGA) of eRF1+eRF3a+GTP and Nsp1 and relative quantitative analysis of the TC formation efficiency on different stop codons (UAA, UAG, UGA) by eRF1+eRF3a+GTP in the presence of Nsp1. TC formation efficiency induced by eRF1+eRF3a+GTP was taken as 1. r.u. – relative units. Positions of preTCs and TCs are labeled by white and black triangles, respectively. Red stars indicate the increased quantity of ribosomal complexes, shifted from the preTC to the TC state. The error bars represent the standard error of mean, stars (**) mark a significant difference from the respective control *P* < 0.01 (n=3).

## DISCUSSION

In the present study, we show that Nsp1 promotes translation termination which is compatible with the finding that Nsp1 binds the 80S ribosome simultaneously with eRF1 and ABCE1 (Thoms et al., 2020). We show the activity of Nsp1 at different stages of translation termination (Fig. 1–3, S1) and identify the amino acid residues of Nsp1 important for its functioning in translation termination (Fig. 4, S2). Additionally, we demonstrate that Nsp1 stimulates termination at all three stop codons (Fig. 6).

We show that Nsp1 stimulates of translation termination on the stage of stop codon recognition because it functions independently of GTP hydrolysis (Fig. 3B) and presence of eRF3 in the reaction (Fig. 1B,2AB). Moreover, it does not stimulate GTPase activity of eRF3, which is stop codon independent (Fig. 3A). Besides analysis of mutant forms of Nsp1 demonstrates that it does not require binding to the mRNA channel of the ribosome to perform its function in translation termination (Fig. 4C). Additionally, we localize interacting regions of these proteins: NM domain of eRF1 and N domain of Nsp1 (Fig. 3C, 4C). Summarizing obtained results, we propose the following molecular mechanism of Nsp1 activity during translation termination – interacting with the NM domain of eRF1 via its N domain, Nsp1 promotes binding of the release factor to the stop codon, which increases the overall efficiency of translation termination.

Nsp1 stimulates expression of host genes via increasing translation termination rate, but at the same time, Nsp1 suppresses cell translation at the initiation stage via binding to the 40S ribosome subunit to transfer components of protein biosynthesis into the translation of virus proteins (Banerjee et al., 2020; Lapointe et al., 2020; Schubert et al., 2020; Shi et al., 2020; Thoms et al., 2020; Tidu et al., 2020; Yuan et al., 2020). That’s why this work could not be done using cell-based experiments, since the effect of Nsp1 on translation termination could not be detected on the background of suppression of translation initiation by this protein. The only possibility to detect its influence on termination in the cells is using two mutants KH/AA and YF/AA which are deficient in translational suppression. These mutants indeed increase the level of the Nluc translation in RRL stimulating peptide release (Fig. 4B).

One can conclude that improving translation termination leads to better gene expression, unless this increased translation termination takes place at sense codons. We tested this possibility and showed that termination is stimulated by Nsp1 specifically at the stop codons (Fig. 6,S5). What are the advantages for the virus to stimulate translation termination? Ribosomes that are in this step will terminate translation and dissociate even without Nsp1, however stimulation of Nsp1 intensifies this process and accelerates cell switching to viral translation. Additionally, Nsp1 probably binds to the mRNA channel at the termination stage to prevent de novo protein synthesis of cellular mRNAs. By stimulating termination in translating mRNA, it can bind to the 40S ribosome subunits and can prevent host initiation of translation. Furthermore, it should be noted that the SARS-CoV-2 mRNA contains two long ORFs for the translation of 16 proteins. They contain only two stop codons, so the elimination of the 40S subunits by Nsp1 after translation termination preferentially involves ribosomes that translate host mRNAs and do not affect the translation of the viral proteins encoded in the first two ORFs. It’s clear that the entire process of translation cannot be stopped by only suppressing initiation because those mRNAs that have already passed the initiation stage continue to be translated at the stages of elongation and termination. To disrupt the translation of such mRNAs, it would be reasonable to stimulate termination at the stop codons and remove the 40S ribosome subunits from the pool of active components. We assume that Nsp1 is involved in this process.

## MATERIAL AND METHODS

### Nsp1 cloning and purification

Codon-optimized Nsp1 wt was amplified by PCR with Q5 polymerase (NEB) from the plasmid pDONR207 SARS-CoV-2 NSP1, which was a gift from Fritz Roth (Addgene plasmid # 141255; http://n2t.net/addgene:141255; RRID:Addgene_141255) (Kim et al., 2020a) using primers NSP1_F, NSP1_R. The SUMO sequence was amplified from the petSUMO plasmid (Invitrogen) by PCR with Q5 polymerase (NEB) using primers petSUMO_F, petSUMO_R. The petSUMO_Nsp1_WT construct was then obtained using PCR-products with Gibson Assembly® Master Mix (NEB), according to the manufacturer’s protocol. N (1-137 aa) and С (139-180 aa) Nsp1 domains were subcloned into petSUMO using primers NSP1_N_R and NSP1_C_F. Nsp1 mutants RK/SE, NK/SE, KH/AA, YF/AA, RR/EE were made with use of QuikChange Site-Directed Mutagenesis Kit (Agilent), according to the manufacturer’s protocol. 3xFLAG was amplified by PCR with Q5 polymerase (NEB) from our construct using primers 3xFLAG_F and 3xFLAG_R and cloned into petSUMO_NSP1_WT vector using Gibson Assembly® Master Mix (NEB). The resulting proteins were expressed in *Escherichia coli* BL21(DE3) cells (Invitrogen) after induction by 0.5 mM isopropyl β-D-1-thiogalactopyranoside at 18°C overnight, followed by purification using a Ni-NTA gravity flow column and elution by 20 mM Tris-HCl pH 7.5, 500 mM KCl, 7,5% glycerol, 300 mM imidazole, and 1 mM DTT. Obtained proteins were dialysed against cleavage buffer (CB), comprising 20 mM Tris-HCl (pH 7.5), 250 mM KCl, 7,5% glycerol, 1 mM DTT. Then peptide bond hydrolysis was carried out with 6xHis-tagged peptidase Ulp1 in CB. Hydrolysis was carried out during the night at +4 º. Then mix was incubated with Ni-NTA agarose, equilibrated in CB, and purified using gravity flow column. The product was dialysed in storage buffer (SB): 20 mM Tris-HCl (pH 7.5), 100 mM KCl, 10% glycerol, 1 mM DTT, frizzed in liquid nitrogen and stored at −70º. The composition of CB for Nsp1 С domain (~4,8 kDa) was identical with SB and the dialysis was not carried out.

### *In vitro* translation assay

The linear template for Nluc were obtained by PCR with the primers: forward primer 5’- CCA GTG CCA AGC TTA ATA CGA CTC ACT ATA G −3’, reverse primer 5’- TTT TTT TTT TTT TTT TTT TTT TTT TTT TTT TTT TTT TTT TTT TTT TTT TTA AAC AGC TAT GAC CAT GAT T −3’. Reverse primers contained poly(T) to generate a polyadenylated 3’end of mRNAs. mRNAs were in vitro transcribed (T7 RiboMAX Express Large Scale RNA Production System, Promega, USA) from the Nluc PCR template. The translation was performed in nuclease-treated RRL obtained as described above. Translation mixture contained 70% nuclease-treated RRL was supplemented with 20 mM Hepes-KOH (pH 7.5), 80 mM KOAc, 0.5 mM Mg(OAc)2, 0.36 mM ATP, 0.2 GTP, 0.05 mM each of 20 amino acids (Promega), 0.5 mM spermidine, 5 ng/μl total deacylated rabbit tRNAs, 10 mM creatine phosphate, 0.003 u/μl creatine kinase (Sigma), 2 mM DTT, 0.2 U/μl Ribolock (ThermoFisher), 1% NLuc substrate (Nano-Glo, Promega), 16 pg/μl NLuc-mRNA and 0.6 μM Nsp1 protein or its mutants/domains or eIF1A as negative control. Luminescence was measured at 30°C during 1 h using a Tecan Infinite 200 Pro (Tecan, Männedorf, Switzerland). The data was presented as luminescence time progress curve with standards deviations, plotted in Microsoft Excel.

### Termi-Luc peptide release assay

A peptide release with Nanoluciferase (NLuc) was performed as previously described (Susorov et al., 2020) with modifications. The assay allows measurement of the NLuc release from TCs assembled on NLuc mRNA in RRL. NLuc folds into the catalytically active form only after its release from the ribosome, which leads to luminescence in the presence of substrate. The NanoLuc mRNA was *in vitro* transcribed (T7 RiboMAX Express Large Scale RNA Production System, Promega, USA) from a template containing β-globin 5’-UTR, NLuc CDS, 3’-UTR derived from pNL1.1[Nluc] vector (Promega), and 50 nt poly(A) tail. 1 ml of RRL lysate (Green Hectares, USA) was preincubated in a mixture containing 1.5 u/μl Micrococcal nuclease (Fermentas) and 1 mM CaCl_2_ at 30°C for 10 min, followed by the addition of EGTA to a final concentration of 2 mM. A reaction mixture containing 70% nuclease-treated RRL was supplemented with 20 mM Hepes-KOH (pH 7.5), 80 mM KOAc, 0.5 mM Mg(OAc)_2_, 0.36 mM ATP, 0.2 GTP, 0.05 mM each of 20 amino acids (Promega), 0.5 mM spermidine, 5 ng/μl total rabbit tRNAs, 10 mM creatine phosphate, 0.003 u/μl creatine kinase (Sigma), 2 mM DTT and 0.2 U/μl Ribolock (ThermoFisher) in a volume of 1 ml. The mixture was preincubated with 1 μM eRF1(AGQ) at 30°C for 10 min, followed by the addition of the NLuc mRNA to a final concentration of 8 μg/ml, resulting in the formation of TCs. Next, KOAc concentration was adjusted to 300 mM and the mixture was layered on 10–35% linear sucrose gradient in buffer containing 50 mM Hepes-KOH, pH 7.5, 7.5 mM Mg(OAc)_2_, 300 mM KOAc, 2 mM DTT. The gradients were centrifuged in a SW-41 Ti (Beckman Coulter) rotor at 18000 rpm for 14 h. Fractions enriched with preTCs were collected. PreTC was aliquoted, flash-frozen in liquid nitrogen, and stored at −80 °C. Peptide release was performed in solution containing 8 nM preTC, 50 mM Hepes-KOH, pH 7.5, 0.25 mM spermidine, 2 mM DTT, 0.2 mM GTP, and 1% NLuc substrate (Nano-Glo, Promega) in the presence of release factors (4 nM eRF1 alone or with 4 nM eRF3a) and 80μM Nsp1 protein or its mutants/domains or eIF1A as negative control. Luminescence was measured for 1 hour at 30°C using a Tecan Infinite 200Pro (Tecan, Männedorf, Switzerland). The peptide release kinetic curves or histogram with maximum values were generated and the standard deviation was calculated in Microsoft Excel.

### Pre-termination complex assembly

The 40S and 60S ribosomal subunits as well as eukaryotic translation factors eIF2, eIF3, eEF1H, and eEF2 were purified from RRL or HeLa lysate, as previously described (Alkalaeva et al., 2006). The human translation factors eIF1, eIF1A, eIF4A, eIF4B, ΔeIF4G, ΔeIF5B, eIF5, PABP, and eRF1 were produced as recombinant proteins in *E. coli* strain BL21(DE3) with subsequent protein purification on Ni-NTA agarose and ion-exchange chromatography (Alkalaeva et al., 2006). The full-length human eRF3a was expressed in insect Sf21 cells and purified via affinity chromatography using a HisTrap HP column (GE Healthcare) followed by anion-exchange chromatography on a MonoQ column (GE Healthcare) (Ivanov et al., 2016). mRNAs were transcribed *in vitro* (T7 RiboMAX Express Large Scale RNA Production System, Promega, USA) from the linear fragment of the pET28-MVHL-UAA plasmid containing the T7 promoter, four CAA repeats, β-globin 5’-UTR, and ORF (coding for the peptide MVHL), followed by the stop codon UAA and 3’-UTR, comprising the remaining natural β-globin coding sequence. Eukaryotic preTCs on MVHL-UAA mRNA were assembled and purified as previously described (Alkalaeva et al., 2006; Egorova et al., 2019; Mikhailova et al., 2017). Briefly, ribosomal complexes were assembled in a 500 μL solution containing 37 pmol of mRNA. They were incubated for 15 min in buffer A (20 mM Tris acetate (pH 7.5), 100 mM KAc, 2.5 mM MgCl_2_, 2 mM DTT) supplemented with 400 U RiboLock RNase inhibitor (Thermo Fisher Scientific, cat# EO0384), 1 mM ATP, 0.25 mM spermidine, 0.2 mM GTP, 75 μg calf liver total tRNA (acylated with all or individual amino acids; Sigma, cat# R7250), 75 pmol human 40S and 60S purified ribosomal subunits, 200 pmol eIF2, 60 pmol eIF3, 400 pmol eIF4A, 150 pmol eIF4B, and 125 pmol each of human eIF2, ΔeIF4G, eIF1, eIF1A, eIF5, and ΔeIF5B for the formation of the 80S initiation complex. After 15 min, 200 pmol of human eEF1H and 38 pmol of eEF2 were added to form the elongation complex. The ribosomal complexes were then purified via centrifugation in a Beckman SW55 rotor for 95 min at 4 °C and 48,000 rpm in a 10– 30% (w/w) linear sucrose density gradient (SDG) prepared in buffer A with 5 mM MgCl_2_. Fractions corresponding to the ribosomal complexes were diluted three-fold with buffer A containing 1.25 mM MgCl_2_ (to a final concentration of 2.5 mM Mg^2+^) and used in experiments.

### Termination complex (TC) formation efficiency assay

First, 0.03 pmol preTCs incubated with release factors (0.5 pmol eRF1/eRF1(AGQ) alone or 0.1 pmol eRF1/eRF1(AGQ) with 0.1 pmol eRF3a) and optionally with 2.5 pmol Nsp1 or its mutants/domains or eIF1A as negative control at 37 °C for 15 min in the presence of 0.2 mM GTP or GDPCP, supplemented with equimolar amounts of MgCl_2_. Samples were analysed using the primer extension protocol. The toe-printing analysis was performed with AMV reverse transcriptase and 5’-carboxyfluorescein-labelled primers complementary to 3’-UTR sequences, as described previously (Egorova et al., 2019). cDNAs were separated via electrophoresis using standard GeneScan® conditions on an ABI Prism® Genetic Analyser 3100 (Applera).

### GTPase assay

Ribosomal 40S and 60S subunits (5 pmol each) with 5 pmol of eRF1 were associated at 37 °C for 10 min. Then, other components were added. Final reaction volume was 10 μL and contained 23.5 mM Tris-HCl pH 7.5, 35 mM NH_4_Cl, 18 mM MgCl_2_, 0.5 mM GTP, 3 pmol of eRF3a and 0, 3, 5, 10 or 15 pmol of Nsp1. After incubation for 15 min at 37 °C the amount of released phosphate was estimated with Malachite Green Phosphate Assay (Sigma), according to the manufacturer’s protocol.

### Quantification and statistical analysis

All experiments were carried out in at least 3 replicates (the specific number of replicates is shown in the description under figures). The data is presented as mean±standard deviation (SD), when analyzing luminescence signals of lysates, or mean±standard error of mean (SE), when analyzing parameters (translation). A two-tailed t-test was used to compare mean values between two groups. For multiple comparisons, the Bonferroni correction was used. One-way ANOVA was used to compare mean values between more than two groups. P-values were calculated in Microsoft Excel. The difference was considered significant when p was less than 0.05 (Glantz, 2011).

## ACKNOWLEDGEMENT

We are grateful to Ludmila Frolova for providing plasmids encoding release factors and to Tatyana Pestova and Christopher Hellen who provided us with plasmids encoding initiation factors. pDONR207 SARS-CoV-2 NSP1 was a kind gift from Fritz Roth (Addgene plasmid # 141255; http://n2t.net/addgene:141255; RRID: Addgene_141255). Sequencing of plasmids, coding mutant proteins and cDNA fragment analyses were performed by the equipment of EIMB RAS “Genome” center (http://www.eimb.ru/ru1/ckp/ccu_genome_c.php).

## AUTHOR CONTRIBUTIONS

A.S., E.S., N.B., E.S., K.E., T.E. and E.A. designed the experiments. A.S., E.S., N.B., E.S., K.E., and T.E. performed the experiments. All authors discussed the results and contributed to the final manuscript. E.A. supervised the project. T.E. and E.A. wrote the manuscript.

## SUPPLEMENTARY DATA

Supplementary Data are available online.

## SUPPLEMENTARY DATA

**Figure S1.** (A) Absence of the TC formation activity of Nsp1 alone. An example of raw toe-printing data. PreTCs are shown after SDG purification. Different amounts of Nsp1 were added to the preTCs, preTC corresponds to white triangle. (B) Saturation curve for eRF3a in the GTPase test. Green arrow shows the concentration of eRF3a used in the following experiments.

**Figure S2.** Activity of the Nsp1 mutants in the TC formation. An example of raw toe-printing data. TC formation was induced by the addition of eRF1 or eRF1+eRF3a+GTP and Nsp1 or five different Nsp1 mutants. An addition of eIF1A was used as a negative control. The TC corresponds to the black triangle, and the preTC corresponds to white triangle. Red stars indicate the increased quantity of ribosomal complexes, shifted from preTC to TC state.

**Figure S3.** Compeering of the activities of untagged Nsp1 and 3xFLAG-Nsp1 in different stages of translation. (A) In vitro Nluc mRNA translation in RRL in presence/absence of 3xFLAG-Nsp1. Time progress curves showing luminescence (in relative luminescence units, rlu), n=3, mean±SD. (B) Termi-Luc peptide release assay in presence/absence of 3xFLAG-Nsp1 and the release factors. Time progress curves showing luminescence (in relative luminescence units, rlu) with NLuc released from ribosome complex upon treatment with the protein of interest, n=3, mean±SD. (C) An example of raw toe-printing data. TC formation was induced by the addition of eRF1 or eRF1+eRF3a+GTP and Nsp1 or 3×FLAG-Nsp1. An addition of eIF1A was used as a negative control. The TC corresponds to the black triangle, and the preTC corresponds to white triangle. Red stars indicate the increased quantity of ribosomal complexes, shifted from preTC to TC state.

**Figure S4.** Activity of the Nsp1 domains in the TC formation. An example of raw toe-printing data. TC formation was induced by the addition of eRF1 or eRF1+eRF3a+GTP and Nsp1 or N/C domains of Nsp1. An addition of eIF1A was used as a negative control. The TC corresponds to the black triangle, and the preTC corresponds to the white triangle. Red stars indicate the increased quantity of ribosomal complexes, shifted from the preTC to the TC state.

**Figure S5.** Lack of the activity of Nsp1 at the sense codons. An example of raw toe print data for ribosomal complexes obtained on the model mRNAs with the sense codons UGG and GCU replacing stop codon. PreTC were obtained by using of charged individual Val, His and Leu tRNAs during elongation, preTC corresponds to white triangle.

