## Supplementary figures and images for "Nsp1 of SARS-CoV-2 Stimulates Host Translation Termination"

### Figure S1

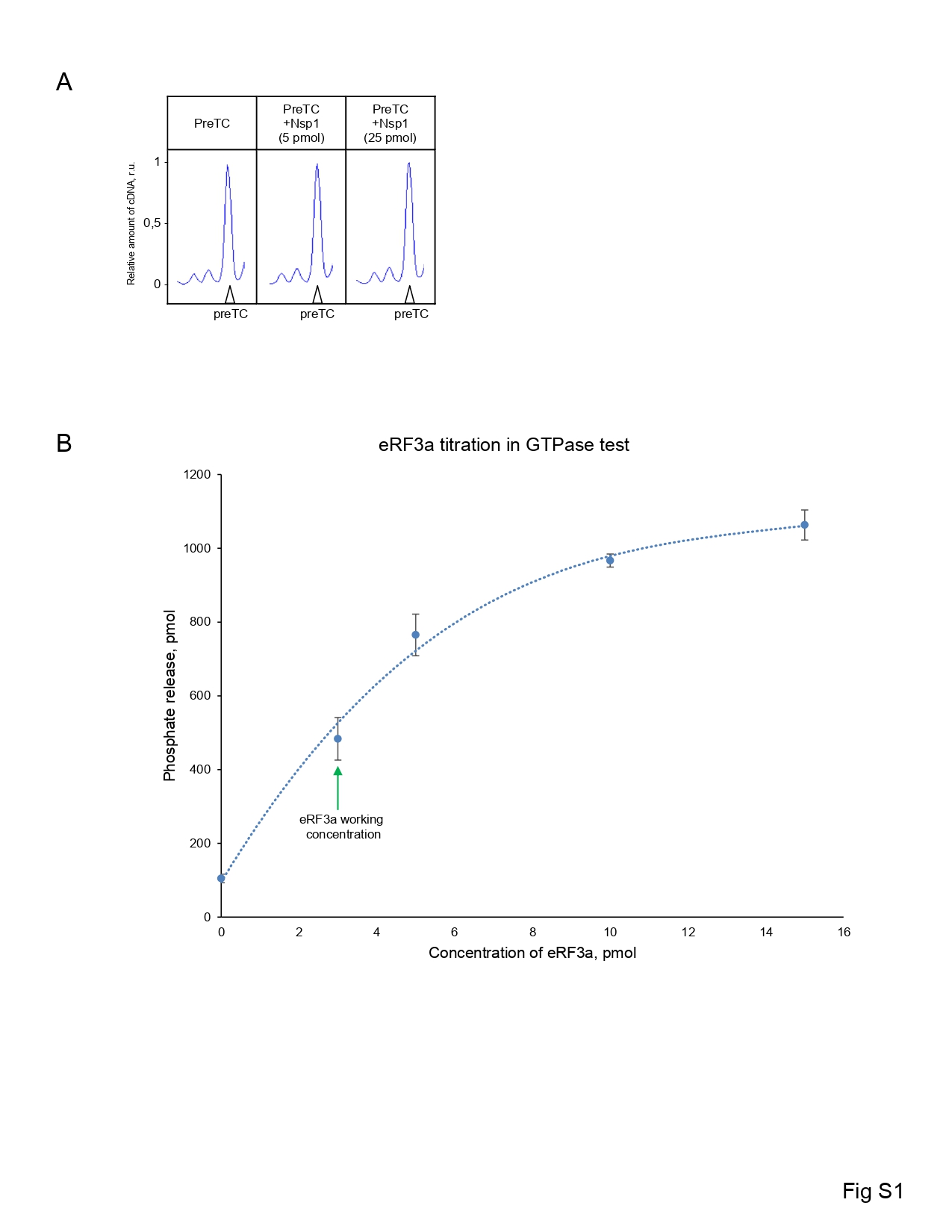

### Figure S2

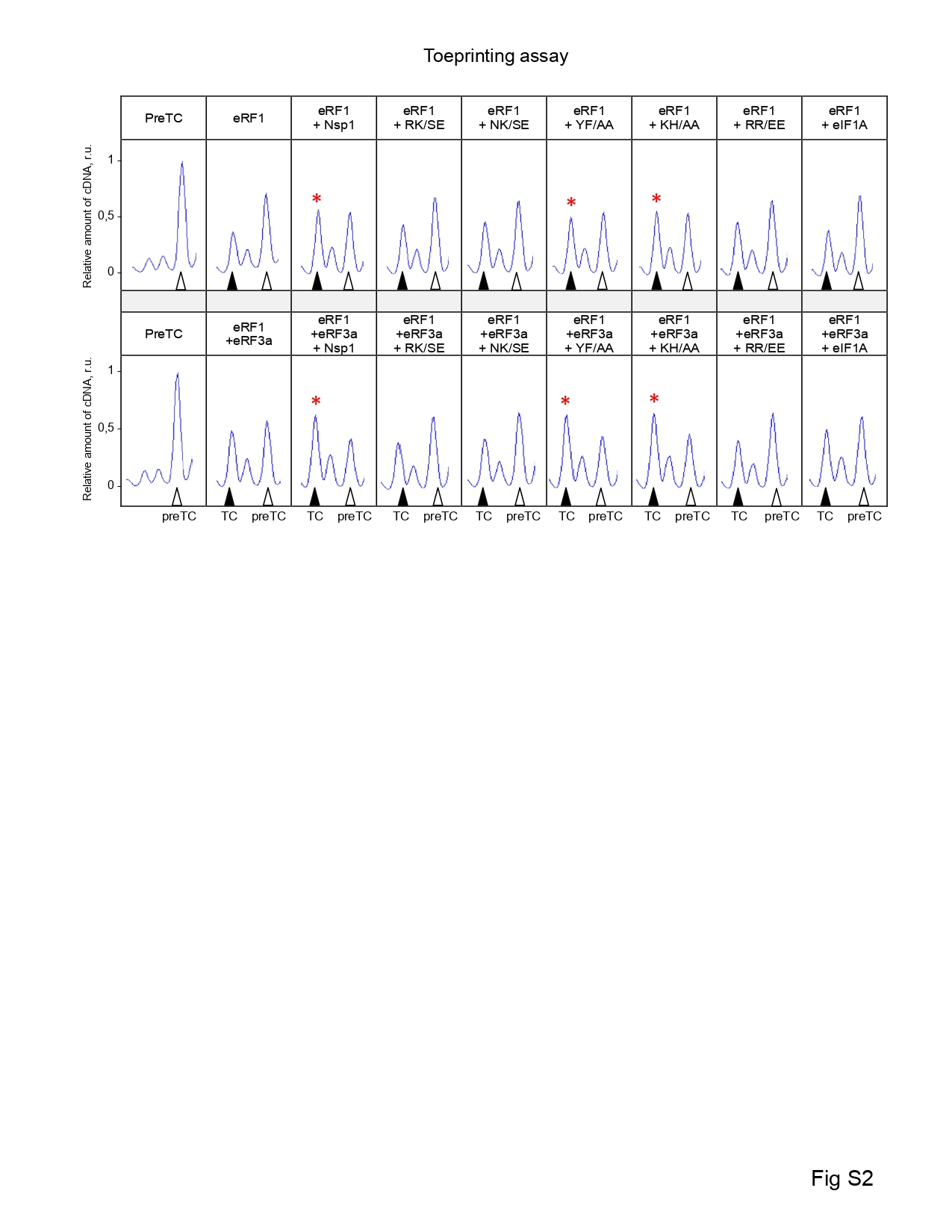

### Figure S3

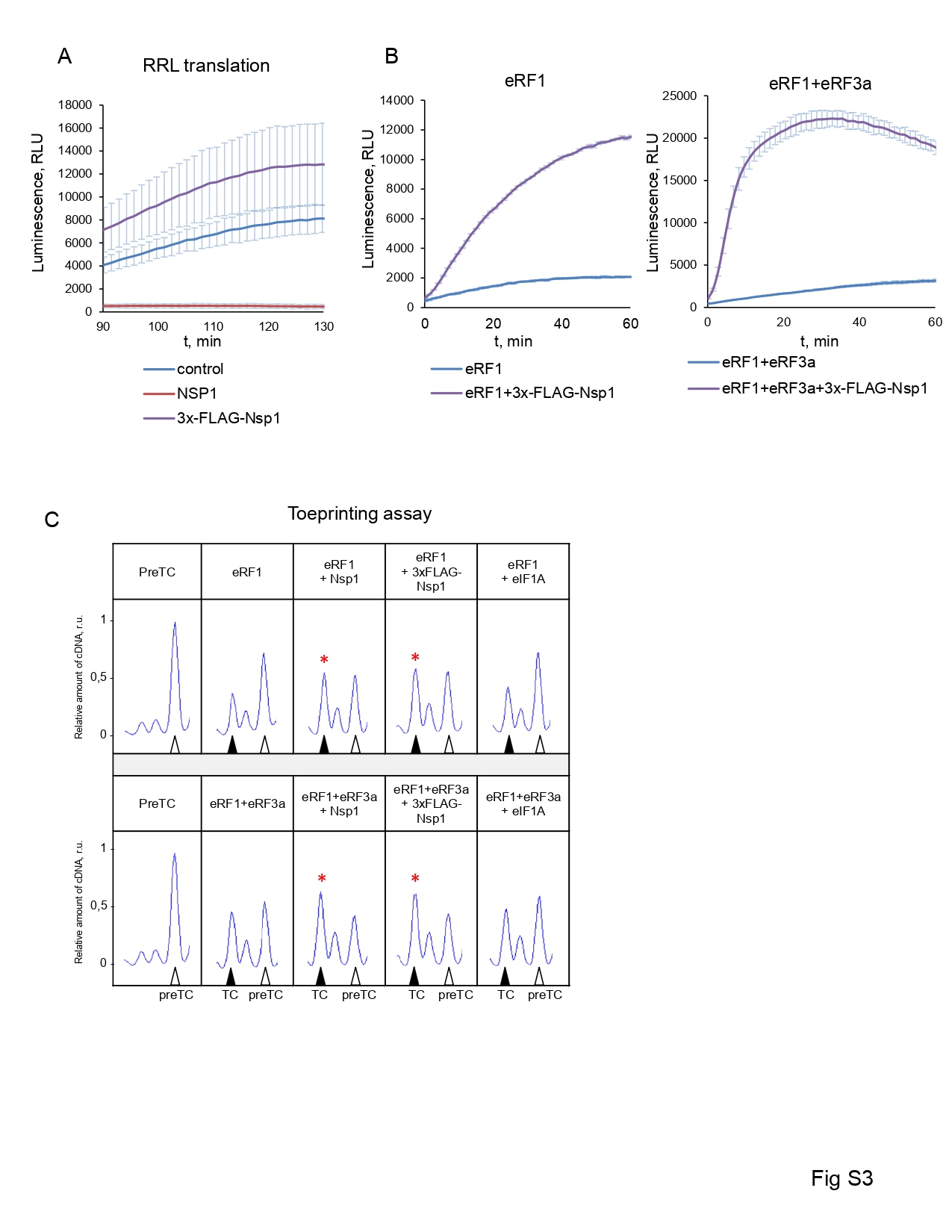

### Figure S4

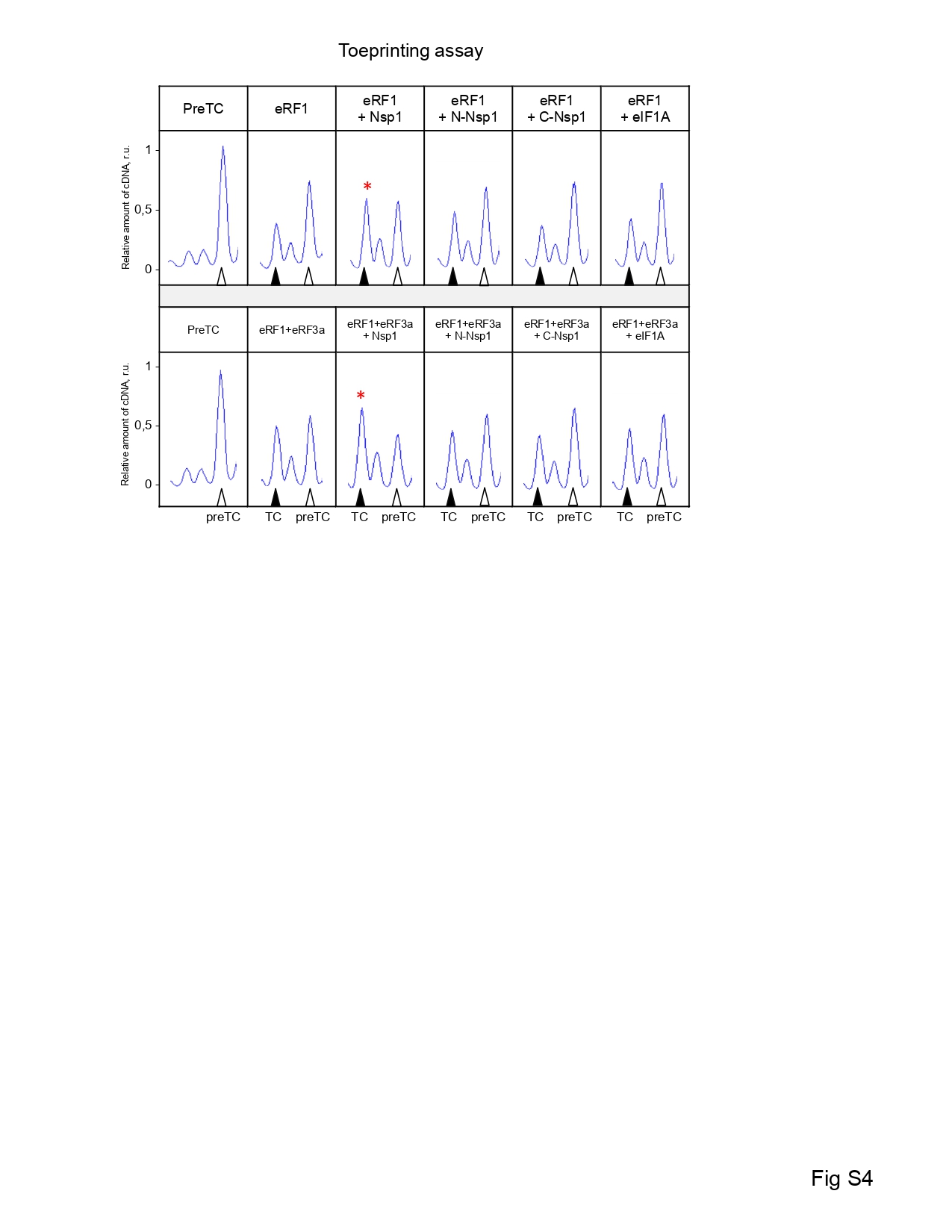

### Figure S5

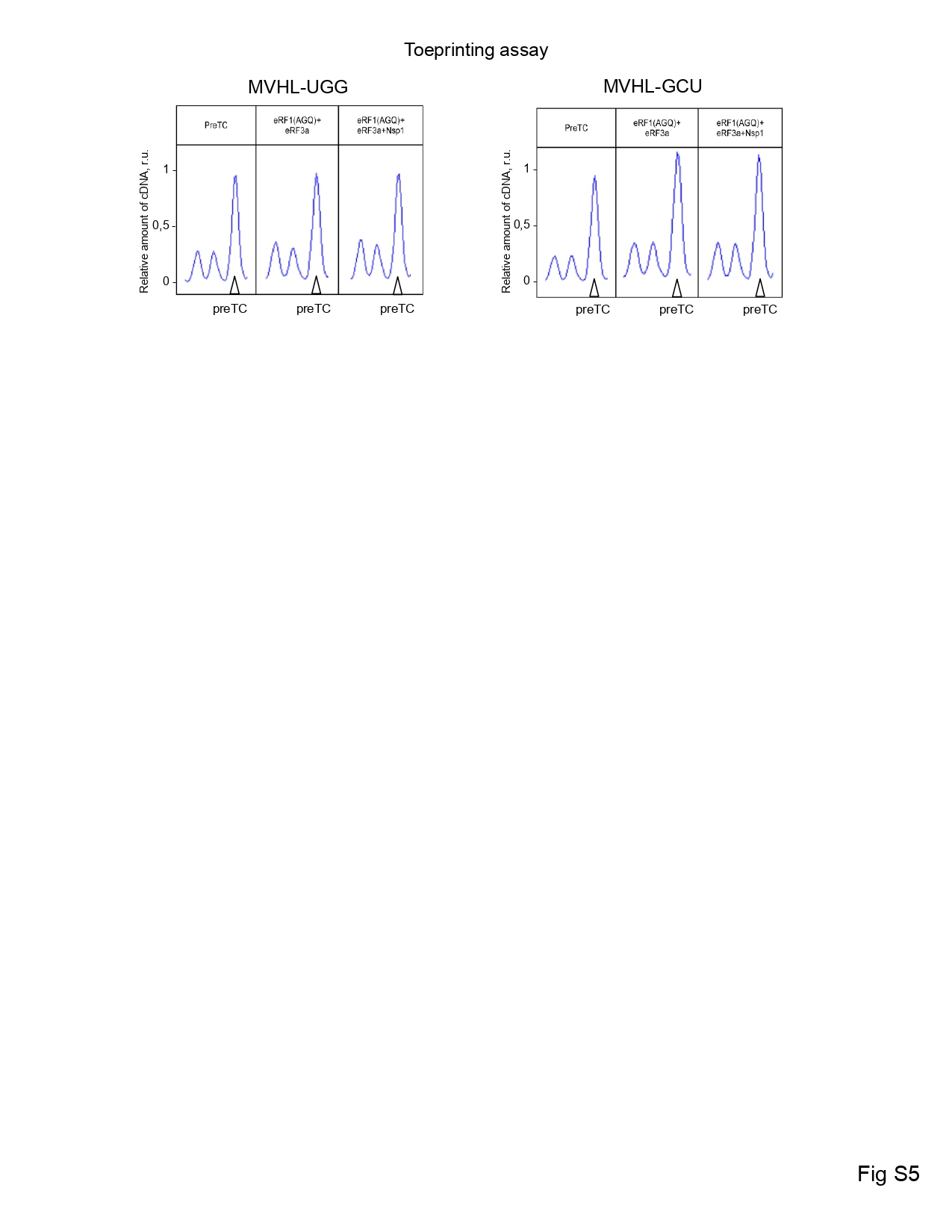
